# Tree growth is better explained by absorptive fine root traits than by transport fine root traits

**DOI:** 10.1101/2024.03.19.585673

**Authors:** Anvar Sanaei, Fons van der Plas, Hongmei Chen, Sophie Davids, Susanne Eckhardt, Justus Hennecke, Anja Kahl, Yasmin Möller, Ronny Richter, Jana Schütze, Christian Wirth, Alexandra Weigelt

**Affiliations:** Institute of Biology, Leipzig University, 04103 Leipzig, Germany; Plant Ecology and Nature Conservation Group, Wageningen University, P.O. Box 47, Wageningen, The Netherlands; Lancaster Environment Centre, Lancaster University, Lancaster LA1 4YQ, UK; German Centre for Integrative Biodiversity Research (iDiv) Halle-Jena-Leipzig, 04103 Leipzig, Germany; Max-Planck-Institute for Biogeochemistry, 07745 Jena, Germany

**Keywords:** absorptive roots, basal area increment, broadleaved tree species, leaves, plant functional traits, root economic space, transport roots

## Abstract

1. Quantifying plant trait variation yields insights into trade-offs inherent in the ecological strategies of plants and is the basis for a trait-based prediction of plant performance and ecosystem functioning. Although the interest in root traits has increased in recent years, we still have limited knowledge of i) whether functionally different fine roots—absorptive versus transport roots—have similar trait coordination and ii) how they help to explain plant performance, such as growth.
2. We measured traits of 25 European broadleaved tree species growing in a research arboretum to study i) the coordination of root traits within absorptive and transport fine roots and ii) the degree of trait-tree growth relationships. To do so, we combined a suite of morphological (root diameter, specific root length and root tissue density) and anatomical (cortex to stele ratio and arbuscular mycorrhizal colonization rate) traits for each of the absorptive and transport roots and also leaf traits (leaf mass per area, dry matter content and toughness).
3. Despite remarkable differences in average trait values between absorptive and transport roots, our study shows that trait coordination within absorptive and transport roots is relatively equivalent. Our results also show that, for the traits we studied, tree growth is better explained by absorptive root traits than by transport root traits and is higher in species with a thinner root diameter. This suggests that variation primarily in absorptive roots affects the uptake of soil-based resources like nutrients and water and directly influences tree growth.
4. The significant relationship between absorptive roots and tree growth and the lack of such a relationship for transport roots highlights that roots mostly involved with resource absorption are more important in explaining tree growth than roots involved in transport.

## 1. Introduction

Functional traits of plants are being used to comprehend plant community structure, assembly and functioning (Lavorel & Grigulis, 2012; Westoby & Wright, 2006). Plant traits reflect different plant strategies and illustrate how plants respond to the environment (Lavorel & Garnier, 2002; Westoby & Wright, 2006); hence, they have the promise to answer how and why plant performance differs among species (Poorter & Bongers, 2006). A suite of associated plant traits known as the leaf economics spectrum (LES) has been established at the leaf level (Reich, 2014; Wright et al., 2004). The LES defines a functional gradient from leaves with conservative resource use to those with an acquisitive strategy, the latter providing a fast return on investment, thus being associated with high growth rates (Reich, 2014). The success of the LES in elucidating variations in leaf traits and predicting plant performance has stimulated researchers to expand the economic theory to ‘fine roots’, proposing a two-dimensional space of roots known as the root economics space (RES; Bergmann et al., 2020). The first dimension is known as the collaboration gradient, and it ranges from species with a high root diameter offering space for arbuscular mycorrhizal fungi to species with a greater specific root length (SRL). The second RES dimension, known as the conservation gradient, is equivalent to the classical LES with high root nitrogen representing a fast-growth strategy and low root tissue density (RTD) representing a slow-growth strategy.

Many ecological studies on root traits define fine roots based on an arbitrary diameter size, and often implicitly assume roots within this size class to be homogenous in their functioning (Pregitzer et al., 2002). However, plant species typically possess hierarchical root systems, so that in reality fine roots are composed of a collection of very heterogeneous orders and branches differing in morphology, architecture, anatomy and longevity (Guo, Li, et al., 2008; Guo, Xia, et al., 2008; McCormack et al., 2015; Pregitzer et al., 2002) as well as in microbial associations (King et al., 2023). Given this, the trait data obtained from different root orders of the same species could be structurally and anatomically different and hence perform different functions (Laliberté, 2017; McCormack et al., 2015). Through this understanding, fine roots have been classified into two distinct groups based on their functional roles. The first group, absorptive roots (order ≤ 3), is responsible for soil-based resource uptake and serves as a hotspot for biotic interactions with microbes and mycorrhizal activity (Freschet & Roumet, 2017; McCormack et al., 2015). The second group, transport roots (order > 3), is most important for transport and storage (Freschet & Roumet, 2017; McCormack et al., 2015). Thus, the capacity of resource transportation increases while absorption capacity decreases with increasing root order (McCormack et al., 2015). Moreover, the lifespan and root diameter of root segments are tied to the location within the branching root system, and consistently increase from the distal to the proximal end (Pregitzer, 2002; Pregitzer et al., 2002). Given this, absorptive roots located at the distal end have a smaller diameter and greater SRL compared to transport roots, and exhibit a shorter lifespan (Pregitzer, 2002; Xia et al., 2010). On the other hand, transport roots, characterised by a larger diameter and longer lifespan, emerge later in the developmental process as a consequence of secondary growth, resulting in greater RTD and lower SRL (Pregitzer, 2002; Xia et al., 2010). In addition, in a root system, anatomical changes across root orders occur mainly due to shifts in physiological functions from resource uptake to transport and storage (Gambetta et al., 2013; Guo, Xia, et al., 2008). As such, a higher percentage of cortex area, or cortex-to-stele ratio, which is characteristic of absorptive roots, is considered an indication of resource absorption and mycorrhizal colonization (Comas et al., 2012; Kong et al., 2017; Zhou et al., 2022). Conversely, a higher stele diameter is known as an indicator of resource transportation in transport roots (Feild & Arens, 2007; Guo, Xia, et al., 2008; Zhou et al., 2022). There is mounting evidence that higher root orders have no cortex due to secondary growth (Endo et al., 2021; Guo, Xia, et al., 2008; Long et al., 2013), thereby reducing mycorrhizal colonization rate (King et al., 2023). Despite this heterogeneity in absorptive and transport root traits, the relative importance of absorptive and transport roots for ecosystem functions such as tree growth is still unexplored.

Forest ecosystem functioning directly and indirectly depends on variation in plant functional traits (Gibert et al., 2016; Paine et al., 2015); thus, studying the link between plant functional traits and ecosystem functioning is important for a mechanistic understanding of forest functioning (Díaz et al., 2016; McGill et al., 2006). Indeed, the effective acquisition and utilization of limited resources are optimized by the functional coordination of roots and leaves and thus their traits (Reich, 2014). Consequently, there has been a lot of interest in identifying the relationship between leaf functional traits and forest functioning (Gibert et al., 2016; Paine et al., 2015; Poorter & Bongers, 2006). For instance, along with the leaf economic spectrum, tree annual growth was positively related to acquisitive traits, characterised by a high specific leaf area (SLA) and stomatal density in subtropical forests (Liu et al., 2015), and a high leaf nitrogen content and SLA in temperate forests (Da et al., 2023). In principle, such relationships have been attributed to higher photosynthetic capacity and a higher potential for a quick return on investment of resources in fast-growing species, leading to a higher growth rate (Reich, 2014; Wright et al., 2004). Even though linking functional traits and plant performance is important, the majority of the studies have reported rather weak links between plant functional traits and plant performance. For example, only 3.1% of variance in tree growth was explained by leaf traits at the global scale in forests (Paine et al., 2015) and 4.8% of variance across functions by leaf and root traits together in grasslands (van der Plas et al., 2020). The reasons for such weak links could be due to the use of species-level mean trait data rather than individual-level trait data and/or using single traits rather than multiple traits, thereby weakening the strength of the relationships between plant functional traits and plant performance. The former might be attributed to the fact that different individuals of the same species respond differently to environmental variables (Siefert et al., 2015); for example, there is some evidence that individual-level trait data improves the degree of trait-growth relationships (Liu et al., 2016; Umaña et al., 2018). Fine roots serve a variety of functions, such as acquiring resources and interacting with soil organisms, all of which influence plant performance (Bardgett et al., 2014; Freschet, Roumet, et al., 2021; McCormack et al., 2015; Smith & Read, 2002). However, our understanding of the relative importance of fine root traits for tree growth lags behind that of leaf traits, partly due to the difficulty of sampling and/or measuring root traits (Freschet, Roumet, et al., 2021). A few recent studies have examined the explanatory power of root traits—in combination with leaf traits—on tree growth, in which for fine roots they focused only on the first three root orders (Shen et al., 2022; Weemstra et al., 2021) or the first two root orders (Da et al., 2023). Shen et al. (2022) showed that acquisitive leaf traits had a higher explanatory power than fine root traits for relative growth rates for height across tree species, even though SRL and RTD were significantly correlated with the relative growth rates for height of individuals. By contrast, Da et al. (2023) found that the conservation gradient of absorptive root traits explained forest aboveground carbon storage and woody biomass productivity better than conservation gradients in leaves and absorptive root collaboration gradients. Although few existing studies have been restricted to absorptive fine root trait effects on tree growth, the simultaneous effects of absorptive and transport root traits have so far been unexplored. Besides, little is known about the effects of anatomical root traits on tree growth. Altogether, this highlights the necessity of examining the trait coordination within functionally discrete fine roots—absorptive and transport roots—as well as examining their relative importance for tree growth, either with or without the combination leaf traits.

By using 25 European broadleaved tree species growing in a research arboretum in Germany, this study aims to quantify the coordination within absorptive and transport fine roots and determine their explanatory power for tree growth, either with or without the combination of leaf traits. More specifically, this study tests the following three hypotheses: First, due to differences in the morphology and anatomical structures between absorptive and transport roots (Guo, Xia, et al., 2008; McCormack et al., 2015; Pregitzer et al., 2002), we hypothesized (H1) that absorptive and transport roots do not necessarily demonstrate similar trait covariation patterns. Second, given the distinct functions of absorptive and transport roots in below-ground processes and functioning (King et al., 2023; McCormack et al., 2015), we hypothesized (H2) that absorptive root traits have a stronger influence on tree growth due to their key role in resource uptake. Third, considering that tree growth relies on concurrent acquisition of above- and below-ground resources, which can be provided through both leaves and roots (Bardgett et al., 2014; Wright et al., 2004), we hypothesized (H3) that tree growth is better explained by a combination of leaf and root traits compared to using root or leaf traits alone.

## 2. Materials and Methods

### 2.1. Study area and experimental design

This study was carried out in the research arboretum ARBOfun located near Leipzig, Germany (51°16′N, 12°30′E; 150 m a.s.l.). The arboretum was established between 2012 and 2014 and is designed for 100 tree species belonging to 39 families planted 5.8 m apart. The 2.5 ha of the arboretum is subdivided into five blocks, with each block containing one individual of each species. The mean annual precipitation is approximately 534.3 mm, and the mean annual temperature is 9.4 °C (Deutscher Wetterdienst (DWD), 2024). The soil type of the arboretum, which was previously used as a managed arable field, is Luvisol, and it has a pH of 5.7 (Ferlian et al., 2017).

### 2.2. Root sampling and measurement

In 2018 and 2019, roots of three individuals per species were sampled. First, the soil around the targeted tree was loosened using a digging fork, and then roots were uncovered carefully by hand and with smaller gardening tools. If a root of higher order was found, it was traced towards the main stem of the target tree to confirm its identity. Then intact root branches containing at least the first five root orders, with the most distal root tip as the first root order, were collected. The root samples, including adherent soil, were wrapped in moist paper, sealed in a plastic bag and stored in a cooling box before being transported to the laboratory. After washing root samples, the sample of each individual tree was divided into two portions: one small portion for examining anatomical traits and another for examining morphological traits. Each subsample comprised fine roots spanning the first to fifth root orders. Finer cleaning was conducted using tweezers under the stereo microscope. After cleaning, the different root orders of the fine root samples were identified and then dissected for trait examination, with each root order being analysed separately. Dissection of root orders was done under a stereo microscope with a scalpel, starting with the root tips as the first root order and categorizing higher root orders towards the stem. From each root sample, 60 root pieces of the first and second root orders, 20 root pieces of the third root order and 10 root pieces of the fourth and fifth root orders were dissected and stored separately in 1.5 ml Eppendorf tubes with water until further processing. The samples of each root order were scanned using a flatbed scanner (Epson Expression 11000XL, UK) at a resolution of 600 dpi, then root pieces were collected, oven-dried at 60°C for over 48 h and weighed to obtain the root dry weight.

All morphological root traits by root orders at individual tree level were quantified using root scans, which were analysed in a batch using the RhizoVision Explorer (Seethepalli et al., 2021). Using the provided data in RhizoVison—mainly the average root diameter, the total root length and volume —alongside the root dry weight data, RTD (root dry weight/root volume) and SRL (total root length/root dry weight) were calculated.

For the measurement of anatomical root traits, root subsamples were cleaned similarly as above, separated by root orders, and placed in scintillation vials containing fixing solution Roti®-Histofix 4% formaldehyde. The samples were left at room temperature for two hours and then refrigerated overnight. The next day, root samples were dehydrated with a series of ethanol with steps of 10%, 30%, 50% and 70%, in which the root samples rested for one hour each to gradually remove the water remained in the root tissue (Zadworny et al., 2016). Samples were kept in the refrigerator in another 70% ethanol solution until further processing. We used an automated tissue processing system (Donatello, Diapath) with (i) 45 min each at 38°C: twice 80% ethanol and twice 96% ethanol, (ii) 60 min each at 38°C and at 40°C xylol and (iii) 80 min each at 62°C three times paraffin, followed by manual embedding of root fragments using a paraffin embedding center (TES 99, Medite). Embedded samples were cross-cut to 1-3 µm with a sledge microtome (DDMP, Medim), put on a slide, processed twice for 10 min in xylol, followed by each 5 min 96%, 80% and 70% ethanol, and finally distilled water before staining for 2 min in 0.01% toluidine blue solution (Aldrich). Slides were permanently fixed with a Tissue Tek system (Sakura). Then, the images of cross-sections per root order were recorded with a microscope (Axiostar plus, Zeiss, Germany) and microscope camera accompanied with the program AxioVision (Zeiss, Oberkochen, Germany). We ensured that the entire cross-section as well as a representative section of higher resolution was depicted in the cross-section image. Analysis of the images for measuring root area, stele area, cortex area and cortex area to stele area ratio (C:S ratio) was done with ImageJ (Schneider et al., 2012).

The rate of arbuscular mycorrhizal colonization (MCR) was investigated using the magnified intersection method (McGonigle et al., 1990). Root pieces were bleached in 10% potassium hydroxide for 18 h. Next, roots were rinsed using deionized water and stained in a 10% ink-vinegar solution (Vierheilig et al., 1998) for 15 min at 90 °C in a water bath. Stained root samples were stored in lactoglycerol until processing. MCR of root pieces was quantified by examining hyphae, arbuscules, hyphal coils, vesicles, and arbuscular mycorrhizal fungi according to the magnified intersection method (McGonigle et al., 1990) with a microscope slide at a magnification of 200x.

According to two distinct groups of fine roots based on their functional roles (McCormack et al., 2015), we used the average of the first three root-order traits to represent absorptive roots and the average of the fourth to fifth root-order traits to represent transport roots for further analyses. In order to verify this classification, we ran additional analyses separately across root orders. The results of these additional analyses are presented in supplementary information (Figure S1), where our main findings and conclusion remain the same, verifying the classification of absorptive and transport roots based on root orders.

### 2.3. Leaf sampling and measurement

13 fully expanded and intact sun-exposed leaves were randomly selected and collected from each individual tree species between 2018 and 2022, following the standard protocol (Cornelissen et al., 2003). Of the 13 leaves, five were scanned at 600 dpi with flatbed Expression 11000XL, and the images were analysed using WinFolia (Regent Instrument, Canada) to get the fresh leaf area. After scanning, the leaf materials were blot-dried, and weighed to get their fresh weights. Then the samples were oven-dried at 60°C for five days and weighed. The leaf mass per area (LMA) was computed by dividing the dry mass of the five leaves (including both lamina and petiole) by their total fresh area. The leaf dry matter content (LDMC) was determined by dividing the mean leaf dry weight by the mean leaf fresh lamina weight. We measured force to punch using a motorised vertical test stand along with a Sauter FH50 with dynamometer combined with a flat-sided needle on three positions of three leaves per species. Additionally, three leaves per species were manually crosscut using a blade to obtain thin sections in the central area of the leaf. The resulting cross sections were then placed in a drop of water on an object slide and examined under a microscope. Then, the mean leaf thickness was determined using the Axiocam (Zeiss, Germany) and the software ZEN 2 core. We then calculated the leaf toughness for each leaf by dividing force to punch of the leaf by the leaf thickness and then computed the individual mean leaf toughness (Westbrook et al., 2011).

### 2.4. Quantification of tree growth

In February 2023, we measured the diameter at breast height (DBH) of each tree individual using a caliper. We then calculated basal area increment as a proxy for tree growth using the sum of DBH data for individual tree species. As such, we calculated the average absolute basal area increment by dividing the 2022 basal area data of each individual tree by its age since planting. Hence, the average basal area increment was calculated according to the following equation:

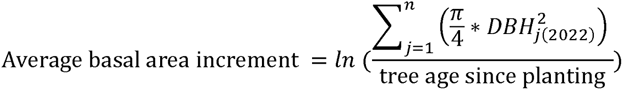

where DBH is the diameter at breath height measured at the 1.3-meter height of an individual, *j* is an index for the *n* stems of the individual, and 2022 is the year when DBH of the individual tree was measured, which overlap the years (2018-2022) during which the trait measurements were done.

### 2.5. Statistical analyses

To assess the variation and coordination of the absorptive and transport root traits, we performed principal component analyses (PCAs) using stepwise inclusion of root and leaf traits at the species average level. To do so, the first set of PCA were performed on morphological and anatomical root traits for absorptive and transport roots separately. The second PCA was performed on leaf traits (LDMC, LMA and LT). Finally, a third set of PCAs were performed on the root traits as well as leaf traits. The PCAs were performed using the *prcomp* function of the ‘stats’ package on scaled trait data and without axis rotation. To aid interpretation, we inverted the PCA axis of the transport root traits by multiplying by minus one whenever required. Next, as the first two PCA axes captured most of the variance, we extracted the loading scores of traits on first and second PCA axes and used them as continuous variables to explain variation in tree growth. Specifically, we performed linear regression to quantify the relationships between average basal area increment (as a dependent variable) and the first and second PCA axes scores (as the explanatory variables) of each PCA coordination using the *lm* function of the ‘stats’ package. Additionally, we used a paired t-test to compare root traits between absorptive and transport roots. To complement the results of PCAs on traits, we subsequently explored the pairwise correlations by performing Pearson’s correlations between absorptive or transport root traits and leaf traits using the *ggraph* function of the ‘ggraph’ package (Pedersen, 2022). To assess each single root and leaf trait as an explanatory predictor for tree growth, we further performed bivariate linear regression separately across absorptive or transport root and leaf traits. To meet the linear regression assumptions, all traits were log-transformed before the regression analysis. All analyses were done using the R v.4.3.2 platform (R Core Team, 2023).

## 3. Results

### 3.1. Covariation in absorptive and transport root traits

We found that root traits, except MCR, substantially differed between different fine root types. In particular, compared to transport roots, absorptive roots showed higher SRL and C:S, while transport roots had higher RD and RTD (Figure 1). Furthermore, pairwise trait correlations showed a much stronger negative correlation between root diameter with RTD and LDMC as well as between MCR and SRL in absorptive roots, while there is a much stronger positive correlation between root diameter with MCR and C:S in absorptive roots (Figure 2).

**Figure 1.**
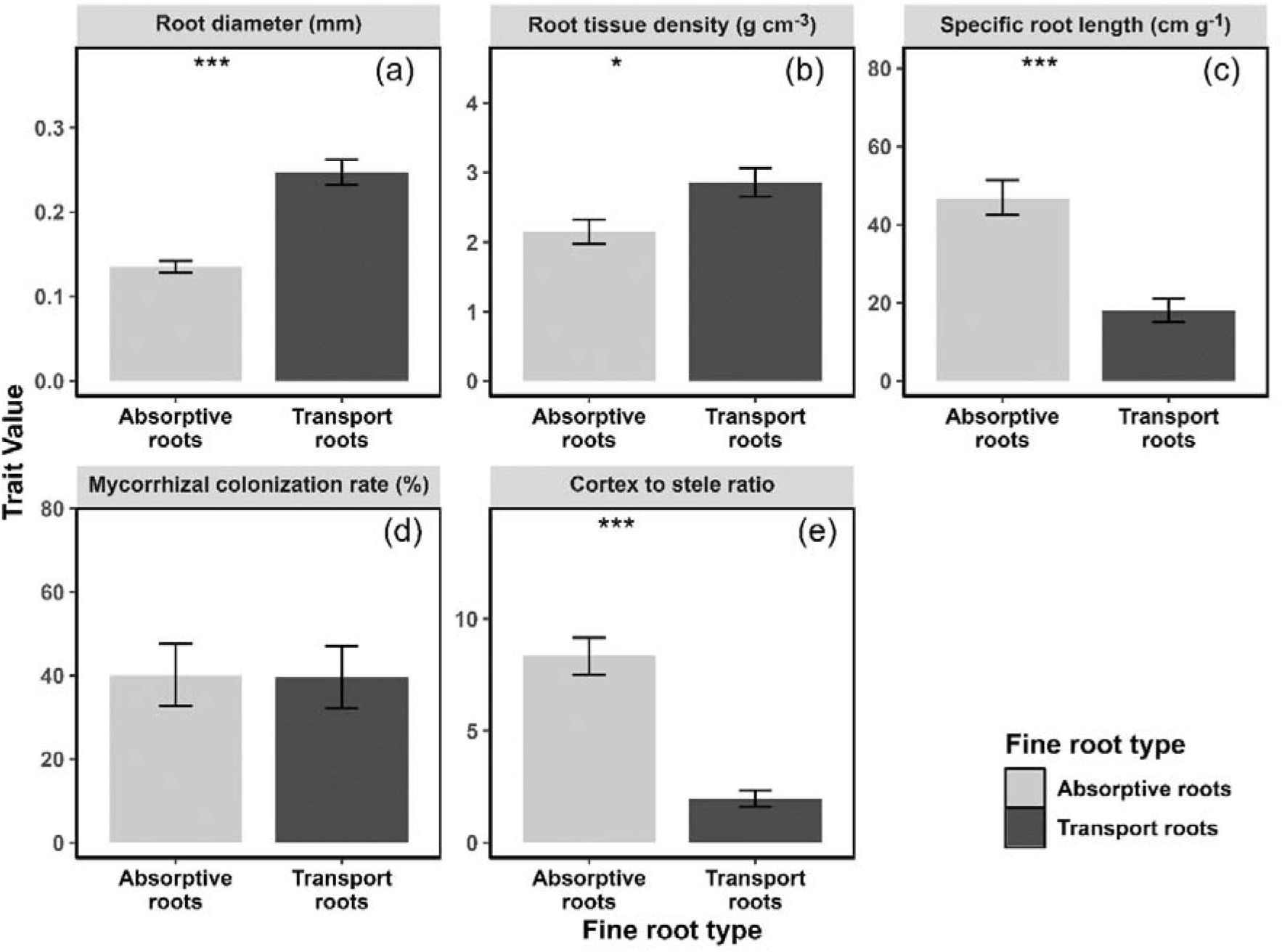
Changes in mean root traits between absorptive and transport roots. Absorptive roots are in light grey, while transport roots are in dark. Significant differences within fine root types are denoted by * (*P < 0.05*), ** (*P < 0.01*) and *** (*P < 0.001*). Data presented are means ± standard error.

**Figure 2.**
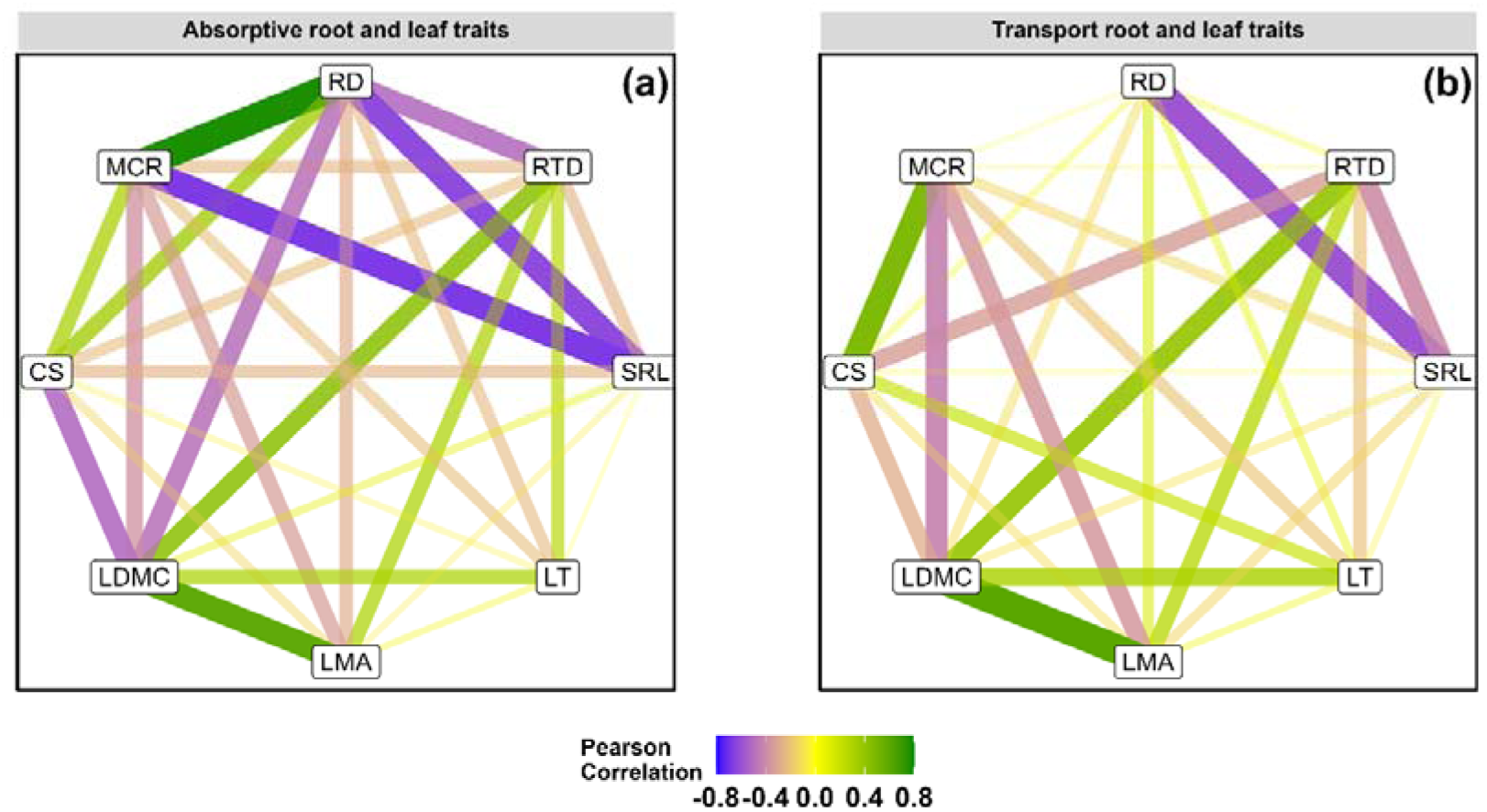
Pairwise trait correlations of (a) absorptive root and leaf traits and (b) transport root and leaf traits. Nodes represent root and leaf traits and line width represents the strength of the correlation. Green and blue lines represent positive and negative correlations, respectively. Abbreviations: RTD, root tissue density; SRL, specific root length; RD, root diameter; CS, cortex to stele ratio and MCR, mycorrhizal colonization rate; LT, leaf toughness; LMA, leaf mass per area; LDMC, leaf dry matter content.

The PCA of absorptive root traits showed that the first two axes together captured 78% of the variability (Figure 3a, Table S2). The first principal component (PCA1) axis is positively associated with root diameter and MCR, and the second principal component (PCA2) axis is positively and negatively related to SRL and RTD, respectively (Figure 3a, Table S2). The first two PCA axes of transport root traits together explained 67% of the variability (Figure 3b, Table S2). The PCA axis 1 of the transport root traits was also positively associated with root diameter, but unlike with absorptive root traits, was in addition negatively associated with SRL (root collaboration gradient), while the PCA axis 2 was positively related to MCR and CS and was negatively related to RTD (Figure 3b, Table S2). Considering only leaf traits, the PCA showed that the first two axes together captured 88% of leaf trait variation (Figure 3c). PCA axis 1 is negatively associated with LDMC and LMA, and PCA axis 2 negatively associated with LT (Figure 3c). The results of the PCA based on the whole set of absorptive root and leaf traits showed that the first two axes accounted for 62% of variation (Figure 3d, Table S2): PCA axis 1 was positively related to the root diameter and MCR while negatively related to LDMC and PCA axis 2 was mainly positively and negatively related to SRL and RTD, respectively (Figure 3d, Table S2). The results based on the whole set of transport root and leaf traits showed that PCA axis 1 and PCA axis 2 accounted for 52% of variation. Unlike with absorptive root traits, PCA axis 1 was negatively related to MCR, while it was positively associated with LDMC and LMA (Figure 3e, Table S2) and PCA axis 2 was negatively associated with SRL, while being positively related to root diameter (Figure 3e, Table S2). While the overall trait coordination of absorptive and transport roots is relatively comparable, MCR and C:S decoupled from RD in transport roots, resulting in MCR and C:S shifting to the second PCA axis in transport roots (Figure 3b).

**Figure 3.**
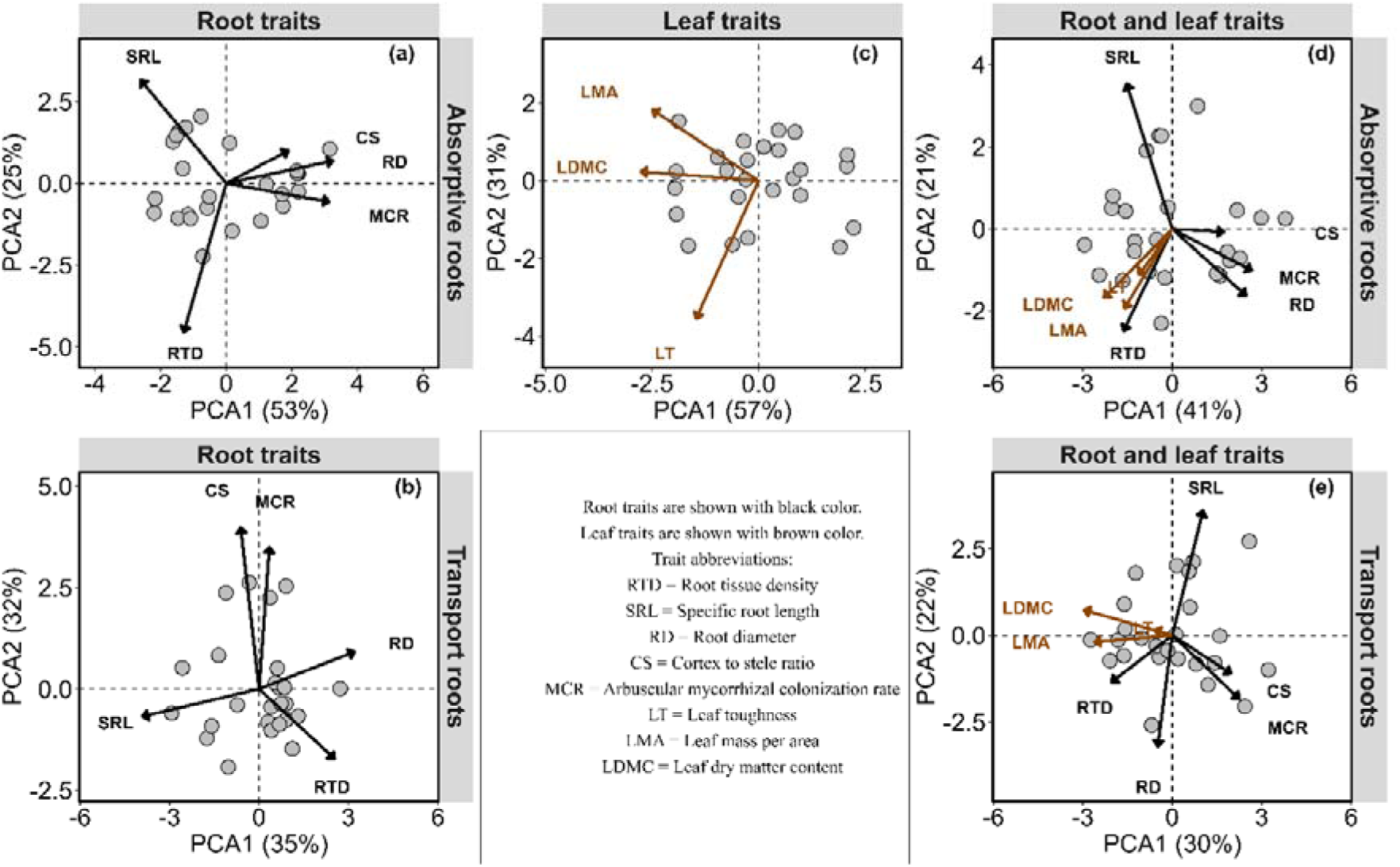
Principal component analyses (PCA) of species-levels of (a,b) absorptive and transport root traits, (c) leaf traits and all root and leaf traits (d,e) for both absorptive and transport root traits. Abbreviations are represented within the figure. The second PCA axis of absorptive morphological traits (d) and the first and second PCA axes of the whole set of transport root and leaf traits (d) are flipped.

### 3.2. The relationships between fine root and leaf traits and tree growth

Our results of linear regressions between PCA axis 1 and average basal tree area increment reveal that absorptive root traits were negatively associated with tree growth (*R^2^* = 0.35, *P* < 0.01; Figure 4a), showing a higher growth for trees with thinner absorptive roots (lower root diameter) and lower MCR, while there was no significant relationship between tree growth and PCA axis 2 (Figure 4b). In contrast, while there was no significant relationship between tree growth and PCA axis 1 of transport root traits (Figure 2d), PCA2 revealed a significant relationship with tree growth (*R^2^* = 0.16, *P* < 0.05; Figure 4d), showing a higher growth for trees with lower C:S and MCR. The linear regressions between PCA axis 1 of leaf traits and average basal tree area increment show that leaf traits were related to tree growth (*R^2^* = 0.20, *P* < 0.05; Figure 4e), showing a higher growth for trees with higher LDMC and LMA. Moreover, PCA axis 1 of absorptive root and leaf traits together explained even more variance in tree growth (*R^2^* = 0.40, *P* < 0.001; Figure 4g), where trees with higher root diameter and MCR but with lower LDMC showed lower growth (Figure 3; Table S2). Finally, PCA axis 1 based on a combination of transport root traits and leaf traits also revealed a significant relationship with tree growth (*R^2^* = 0.18, *P* < 0.05; Figure 4i), where trees with lower MCR but with higher LDMC and LMA showed higher growth (Figure 4i; Table S2). The explanatory power of absorptive root and leaf traits on tree growth was much stronger (Figure 4g) than that of transport root and leaf traits (Figure 4i).

**Figure 4.**
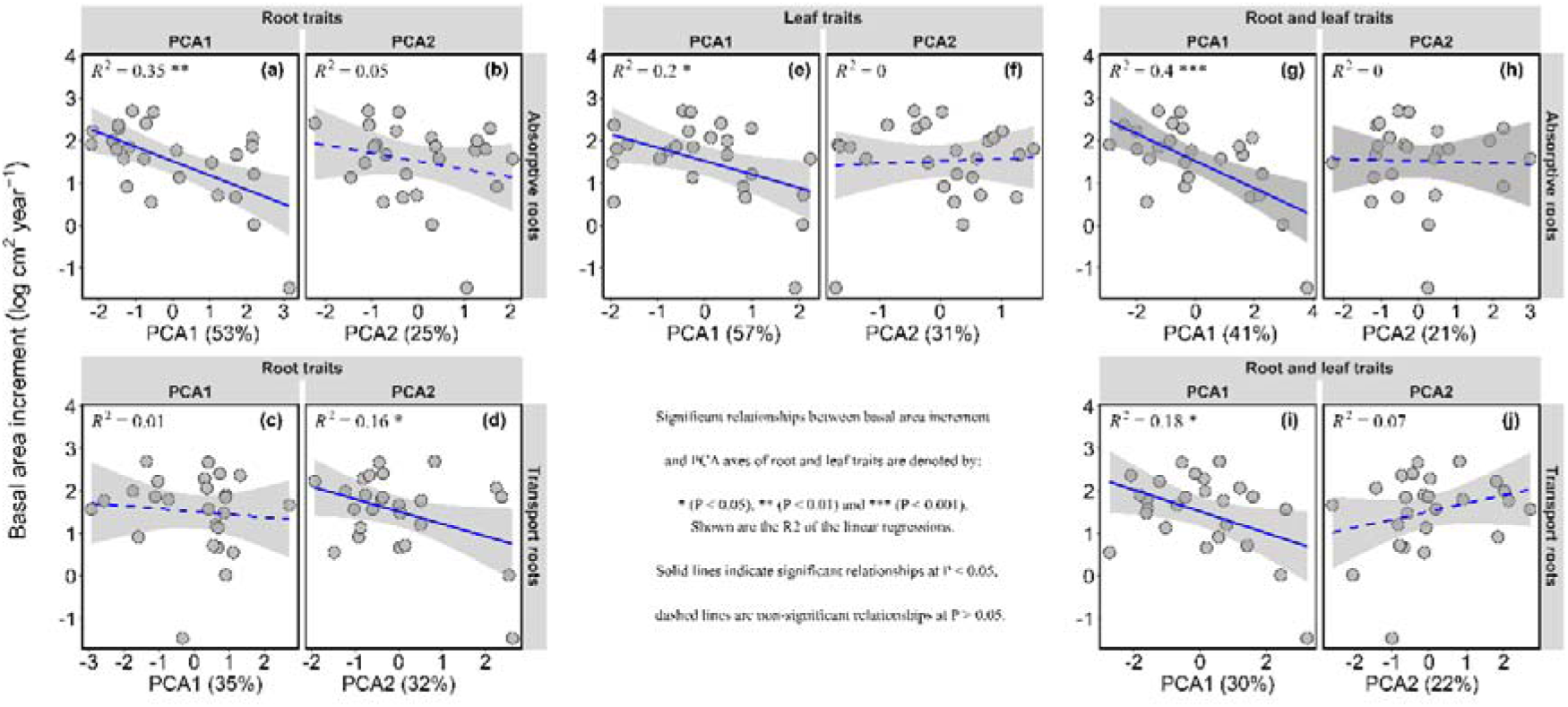
Relationships between the first and second axis of the PCA and average basal area increment and (a,b) absorptive root traits, (c,d) transport root traits, (e,f) leaf traits and (g,h) absorptive and (i,j) transport root and leaf traits. Shown are the *R*^2^ and *P*-values of the linear regressions. Significant relationships between basal area increment and PCA axes are denoted by * (*P < 0.05*), ** (*P < 0.01*) and *** (*P < 0.001*).

## 4. Discussion

By functionally separating fine roots into absorptive and transport roots and also by stepwise inclusion of root traits in PCA, we explored the coordination within absorptive and transport fine roots, which, based on our knowledge, has not been tested so far. Overall, we found that trait coordination within absorptive and transport roots is comparable. Specifically, mycorrhizal colonization, root diameter, and cortex-to-stele ratio were the key traits loading on the first PCA axis. Furthermore, tree growth is better explained by absorptive root traits than by transport roots and was higher in species with thinner root diameter that were less colonized by arbuscular mycorrhizae, highlighting the role of efficient and independent exploration of soil resources.

### 4.1. Covariation in absorptive and transport root traits

Despite significant differences between absorptive and transport root traits (Figure S1), we found that, contrary to our first hypothesis (H1), coordination within absorptive and transport root traits was quite similar to each other and similar to the collaboration gradient published previously; whereby species with higher root diameter were highly related to mycorrhizal association (Bergmann et al., 2020). In partial disagreement with our results, in another study different economic strategies were observed for thin (<L247Lµm) and thick (>L247Lµm) fine roots, where thin roots followed the resource acquisition-conservation strategy but thick roots did not (Kong et al., 2016). It must be mentioned that Kong et al. (2016) applied univariate regression analysis between root traits, not PCA for the trait coordination. The specific fine root diameter cutoff, limited number of species, and/or inclusion of root nitrogen concentration, which we did not measure, can contribute to the different observed patterns. This again highlights the importance of trait selection for the outcome of studies on trait coordination patterns (Weigelt et al., 2023).

Against our expectation, there was no significant difference in mycorrhizal colonization between absorptive and transport roots (Figure S1), which is contrary to the generally acknowledged notion that higher root orders (or transport roots) are not or less colonized by mycorrhizae (King et al., 2023; McCormack et al., 2015). Indeed, transport roots possess lower potential for mycorrhizal colonization due to their thinner cortex (or presence of periderm), providing smaller space for mycorrhizal colonization (Eissenstat et al., 2015; Kong et al., 2017; McCormack et al., 2015). Our inconsistent results might be partly attributed to topological root order classification, in which a higher proportion of thinner roots (absorptive roots) are classified as having higher root orders (transport roots); therefore, the percentage of transport roots is higher than in the morphometric root order classification method (Freschet, Pagès, et al., 2021). Moreover, species-specific differences in mycorrhizal dependence might affect the overall colonization of the roots with mycorrhizae (Zhou et al., 2022). There is some evidence that, for example, *Fraxinus rhynchophylla* Hance. has mycorrhizal colonization in fourth order roots and *Acacia auriculiformis* A.Cunn. ex Benth. is colonized even in fifth order roots, meaning that some species are more colonized by mycorrhizae than others even in higher root orders (Guo, Xia, et al., 2008; Long et al., 2013). This is because plant species differ in the secondary growth development, and mycorrhizal colonization in higher root orders also confirms a higher dependency of those species on mycorrhizae for nutrient uptake (Zhou et al., 2022). This was the case in our mycorrhizal colonization data. As such, order-based root mycorrhizal colonization data showed that for the majority of species, mycorrhizal colonization was greater in the lower root orders or remained on the same level in the higher root orders. Yet, in some species, like *Fraxinus excelsior* L., *Euonymus europaeus* L. and *Frangula alnus* L. mycorrhizal colonization slightly increased with increasing root orders. Altogether, this might lessen the overall mycorrhizal colonization rate differences between absorptive and transport roots.

By incorporating leaf traits into PCAs with absorptive and transport roots, trait coordination showed that conservative leaf traits were closely aligned with conservative root traits, reaffirming that the conservation gradients of both leaf economic spectrum and root economic space are correlated (Reich, 2014). Similar results have been reported when leaf and root traits were pooled, indicating the same trade-offs between the fast–slow conservation gradient in root and leaf traits (Kramer-Walter et al., 2016; Weigelt et al., 2021).

### 4.2. Absorptive root traits better explain tree growth than transport root traits

Past attempts at exploring the contribution of fine root traits to plant performance have considered fine roots as a homogenous pool without regard to their distinct functional roles (Orwin et al., 2018; van der Plas et al., 2020). Thus far, our understanding of how fine roots contribute to tree growth stems from studies testing either the first two or three root orders (Da et al., 2023; Shen et al., 2022; Weemstra et al., 2021), but there is no study testing the effects of functionally discrete fine roots on tree growth. By separating fine roots into absorptive and transport roots, we found that absorptive fine root traits are highly correlated with tree growth, consistent with our second hypothesis (H2). The greater contribution of absorptive root traits to tree growth compared to transport root traits can be attributed to the functioning role of absorptive roots within the plant system (Freschet & Roumet, 2017; McCormack et al., 2015). Within the plant, absorptive roots are mainly involved in soil-based resource acquisition (e.g., nutrients and water), which is directly linked to tree growth. More specifically, the absorptive root traits loaded on the PCA axis 1 (MCR, root diameter and C:S ratio) were the key traits associated with tree growth, highlighting the importance of thin roots with a ‘do-it-yourself’ strategy of resource uptake for tree growth (Lynch et al., 2021; Bergmann et al., 2020). Indeed, the positive associations among mycorrhizal colonization, root diameter and cortex-to-stele ratio are characteristic of absorptive roots (Smith & Read, 2002), and we observed that those traits have stronger correlations in absorptive roots (Figure 2). More precisely, our results showed that species with thicker roots that are more colonised by arbuscular mycorrhizal fungi (Comas et al., 2012; Eissenstat et al., 2015) were negatively correlated with tree growth. Indeed, plants with thicker roots tend to have a longer lifespan and a smaller surface area, resulting in a smaller volume of below-ground resources explored and thus a high dependence on mycorrhizal colonization (McCormack & Iversen, 2019; Pregitzer et al., 2002). In contrast, SRL, as a part of root collaboration gradient in root economic space, was positively correlated with tree growth, meaning that species with the ability to independently explore soil for resources have a higher growth rate. Similar results were obtained based on single-trait bivariate relationships, where root diameter, and cortex-to-stele ratio were significantly and negatively correlated with tree growth and mycorrhizal colonization rate was marginally negatively significant, while SRL and RTD were marginally positively correlated with tree growth (Figure S2). In addition, tree species in Leipzig were experiencing a drought from 2018 to 2020 (Schnabel et al., 2022), so it seems that thinner and smaller root diameters, i.e., potentially reaching smaller pores of soil, are more beneficial, particularly during dry years (Comas et al., 2013), thereby enabling the acquisition of higher nutrients and water with low investment. A higher SRL and smaller root diameter are associated with higher hydraulic conductivity, which reflects drought tolerance capacity (Comas et al., 2012, 2013; Hernández et al., 2010).

Leaf traits were significantly related to tree growth, both alone and when combined with absorptive roots, in support of our third hypothesis (H3), showing that tree growth is best explained by a combination of leaf and root traits. Leaves play crucial roles in plants by converting sunlight energy, carbon dioxide and water into organic carbon through photosynthesis (Reich, 2014; Schulze, 2006; Wright et al., 2004), thereby influencing growth. Our results corroborate previous studies showing the importance of leaf traits and their contribution to forest functioning (J. Cornelissen et al., 1996; Poorter & Bongers, 2006). Our results showed that thinner absorptive roots that are less colonized with arbuscular mycorrhiza were related to high tree growth, but this was the case only for species with higher LDMC and LMA. More specifically, species with a slow-growing strategy (higher LDMC and LMA) tend to have higher growth, indicating a decoupled root and leaf trait strategy explaining tree growth. The opposite pattern has been reported, where species with a high root diameter but a lower specific leaf area enhanced tree growth (Weemstra et al., 2021). Taken together, these findings confirm that traits more directly related to resource uptake above- and below-ground are important indicators of tree growth (Weemstra et al., 2021; Weigelt et al., 2021).

While the second PCA axis of transport roots, indicating species with a higher cortex-to-stele ratio that are more colonized with arbuscular mycorrhiza, is significantly related to tree growth, its predictive power is only half as strong as that of transport roots (Figure 4a,d). We believe that the significant relationship between transport roots and tree growth might be related to similar mycorrhizal colonization rates of absorptive and transport roots (Figure 4a,d). In addition, incorporation of leaf traits to transport root traits slightly increased the explanatory power of estimating tree growth. Based on the trait loading on the PCA axes (Table S2), we believe that this contribution arises from leaf traits rather than transport root traits, as LDMC and LMA are primarily loaded on the first PCA axis (Figure 1f, Table S2). The smaller explanatory power of transport roots in tree growth compared to absorptive roots confirms that transport roots are mainly involved in the transport and storage of resources and also play crucial roles in protecting plants against pathogens and dehydration (Enstone et al., 2002; Lynch et al., 2021) rather than resource acquisitions that are directly related to growth (McCormack et al., 2015).

By functionally separating fine roots into absorptive and transport roots, our results show a strong association between absorptive fine root traits and broadleaved tree growth in a research arboretum. A higher contribution of absorptive root traits to predicting tree growth suggests that variation in absorptive root traits, rather than transport root traits, better explains tree growth variation via presumably providing soil-based resources, e.g., nutrients and water, directly influencing overall tree growth. We argue that by considering fine roots (≤ 2 mm in diameter) as a homogenous pool, the variance of root traits along root orders might be underestimated and might not clearly show root functioning signals. We also acknowledge that further research assessing the role of root and leaf nutrient concentrations as well as considering transport root-related functions may be particularly illuminating.

## Acknowledgments

We thank Roman Patzak, Imke Pelloth, Lea von Sivers, Tom Künne and Julia Leonore van Braak for their help with the field and lab measurements. We also thank Maritta Wipplinger from the institute of veterinary pathology at Leipzig University for her help in root cross-section preparation. We are especially grateful to Florian Schnabel, Lena Kretz and David Schellenberger Costa for helping with discussing ideas. A.S is supported by the Saxon State Ministry for Science, Culture and Tourism (SMWK) – [3-7304/35/6-2021/48880].

## Author Contributions

A.W., C.W. and A.S. conceived the ideas and developed the concept of the study. F.v.d.P., H.C., S.D., S.E., A.K., J.M., J.S. and A.W. contributed to data collection. A.S. analysed the data and led the writing of the manuscript. F.v.d.P., H.C., R.R., J.H., C.W. and A.W. contributed to the writing in several manuscript interactions. All authors contributed critically to the drafts and gave final approval for publication.

## Data Availability Statement

We will store the dataset in the public repository once the paper is accepted.

