## Supporting Information for "Tree growth is better explained by absorptive fine root traits than by transport fine root traits"

**Table S1.** Species information in the study.

| Species | Genus | Family |
|---|---|---|
| *Acer platanoides* | Acer | Sapindaceae |
| *Acer platanoides* | Acer | Sapindaceae |
| *Aesculus hippocastanum* | Aesculus | Sapindaceae |
| *Alnus glutinosa* | Alnus | Betulaceae |
| *Alnus incana* | Alnus | Betulaceae |
| *Betula pubescens* | Betula | Betulaceae |
| *Castanea sativa* | Castanea | Fagaceae |
| *Corylus avellana* | Corylus | Corylaceae |
| *Euonymus europaeus* | Euonymus | Celastraceae |
| *Fagus sylvatica* | Fagus | Fagaceae |
| *Frangula alnus* L. | Frangula | Rhamnaceae |
| *Fraxinus excelsior* | Fraxinus | Oleaceae |
| *Fraxinus ornus* | Fraxinus | Oleaceae |
| *Juglans nigra* | Juglans | Juglandaceae |
| *Mespilus germanica* | Mespilus | Rosaceae |
| *Ostrya carpinifolia* | Ostrya | Corylaceae |
| *Prunus mahaleb* | Prunus | Rosaceae |
| *Quercus cerris* | Quercus | Fagaceae |
| *Quercus robur* | Quercus | Fagaceae |
| *Quercus rubra* | Quercus | Fagaceae |
| *Salix alba* | Salix | Salicaceae |
| *Salix pentandra* | Salix | Salicaceae |
| *Sorbus aucuparia* | Sorbus | Rosaceae |
| *Sorbus torminalis* | Sorbus | Rosaceae |
| *Ulmus laevis* | Ulmus | Ulmaceae |

**Table S2.** Results of the principal component analysis based on the correlation matrix of 25 broadleaves tree species for fine root (absorptive and transport roots) and leaf traits as shown in Figure 3. SRL is specific root length; RTD is root tissue density; RD is average root diameter; C:S is cortex to stele ratio; MCR is mycorrhizal colonization rate; LDMC is leaf dry matter content; LMA is leaf mass per area; and LT is leaf toughness. Displayed data are the variance explained by each principal component and the loading scores of the root and leaf traits on the first four PCA axes.

| | *Absorptive root traits* | | | | | *Transport root traits* | | | |
|---|---|---|---|---|---|---|---|---|---|
| **Fig. 3a** | **PCA1** | **PCA2** | **PCA3** | **PCA4** | **Fig. 3b** | **PCA1** | **PCA2** | **PCA3** | **PCA4** |
| Variance | 0.531 | 0.255 | 0.156 | 0.047 | Variance | 0.348 | 0.317 | 0.193 | 0.082 |
| SRL | -0.46 | 0.55 | 0.03 | 0.37 | | -0.69 | -0.11 | -0.02 | -0.10 |
| RTD | -0.23 | -0.80 | 0.21 | 0.10 | | 0.44 | -0.30 | -0.66 | 0.33 |
| RD | 0.57 | 0.12 | -0.24 | -0.45 | | 0.56 | 0.16 | 0.56 | -0.19 |
| MCR | 0.55 | -0.10 | -0.19 | 0.81 | | 0.06 | 0.61 | -0.50 | -0.60 |
| C:S | 0.33 | 0.18 | 0.93 | 0.02 | | -0.11 | 0.70 | 0.03 | 0.70 |

| | *Leaf traits* | | |
|---|---|---|---|
| **Fig. 3c** | **PCA1** | **PCA2** | **PCA3** |
| Variance | 0.571 | 0.314 | 0.115 |
| LDMC | -0.69 | 0.06 | 0.72 |
| LMA | -0.62 | 0.46 | -0.63 |
| LT | -0.37 | -0.89 | -0.28 |

| | *Absorptive root and leaf traits* | | | | | *Transport root and leaf traits* | | | |
|---|---|---|---|---|---|---|---|---|---|
| **Fig. 3d** | **PCA1** | **PCA2** | **PCA3** | **PCA4** | **Fig. 3e** | **PCA1** | **PCA2** | **PCA3** | **PCA4** |
| Variance | 0.412 | 0.208 | 0.123 | 0.104 | Variance | 0.298 | 0.216 | 0.175 | 0.123 |
| SRL | -0.28 | 0.64 | 0.01 | 0.04 | | -0.19 | -0.66 | 0.00 | 0.05 |
| RTD | -0.30 | -0.45 | -0.22 | -0.01 | | 0.37 | 0.25 | -0.44 | 0.35 |
| RD | 0.48 | -0.18 | 0.14 | -0.13 | | 0.09 | 0.58 | 0.19 | -0.48 |
| MCR | 0.45 | -0.30 | 0.07 | -0.24 | | -0.41 | 0.33 | -0.15 | -0.51 |
| C:S | 0.31 | -0.01 | -0.35 | 0.78 | | -0.36 | 0.20 | 0.42 | 0.47 |
| LDMC | -0.42 | -0.31 | 0.22 | -0.13 | | 0.54 | -0.13 | 0.17 | 0.35 |
| LMA | -0.29 | -0.35 | 0.41 | 0.52 | | 0.47 | 0.03 | 0.15 | 0.22 |
| LT | -0.20 | -0.20 | -0.77 | -0.16 | | 0.11 | 0.03 | 0.72 | 0.04 |

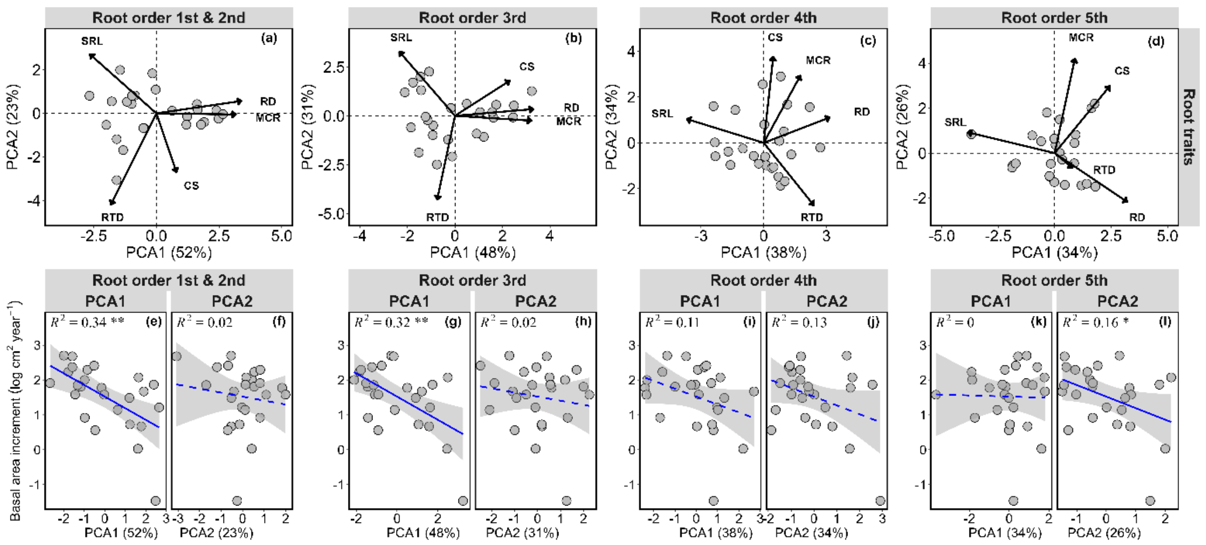


**Figure S1.** Principal component analyses (PCA) of root traits across root orders (a-d), and the relationships between the PCA axes of root traits and average basal area increment (e-l). Shown are the $R^2$ of the linear regressions. Significant relationships between basal area increment and PCA axes are denoted by * (*P < 0.05*), ** (*P < 0.01*) and *** (*P < 0.001*). Abbreviations for traits are as follows: RD, root diameter; RTD, root tissue density; SRL, specific root length; C:S, cortex to stele ratio; MCR, mycorrhizal colonization rate; LMA, leaf mass per area; LT, leaf toughness; LDMC, leaf dry matter content.

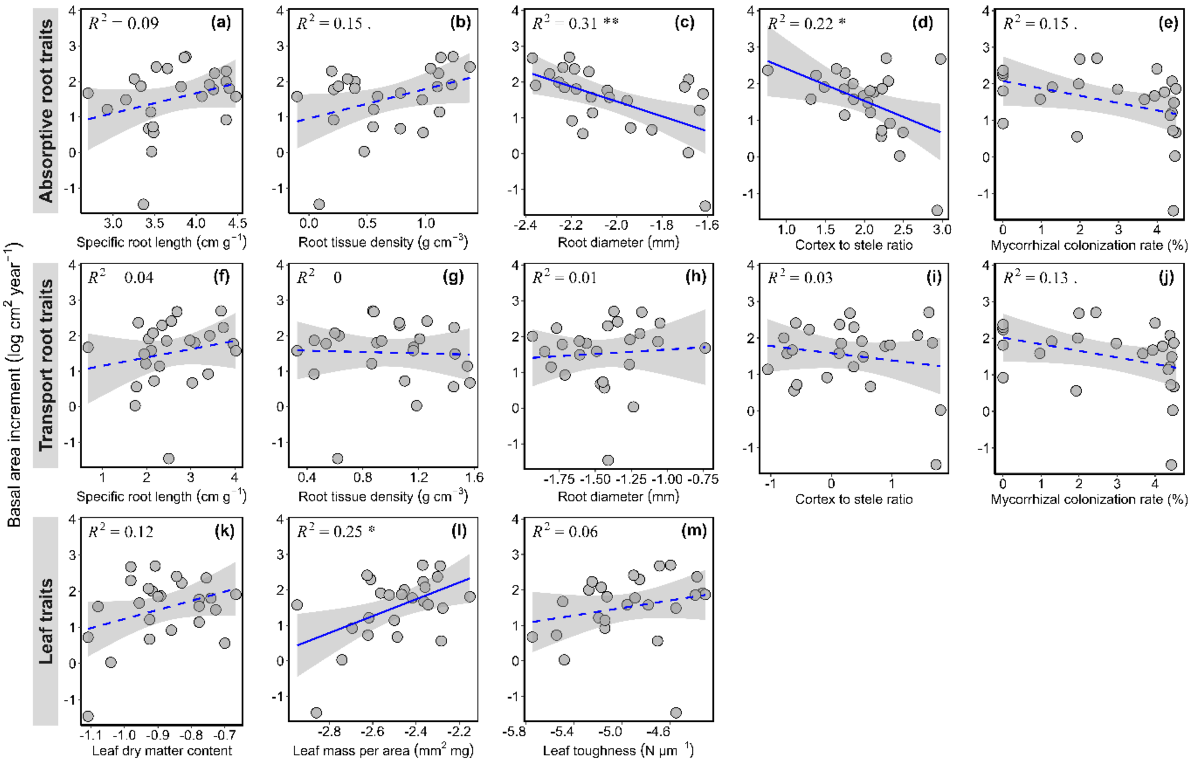


**Figure S2.** The relationships between a:e) absorptive root traits, and f:j) transport root traits, and k:m) leaf traits with average basal area increment. Log-transform data is used in regression. Significant correlations are indicated as solid lines and non-significant relationships as dashed lines, and the shaded areas denote the 95% confidence interval. Shown are the $R^2$ of the regressions, which refers to the proportion of variance in basal area increment explained by root and leaf traits. Significant relationships between basal area increment and root or leaf traits are denoted by * (*P < 0.05*), ** (*P < 0.01*) and *** (*P < 0.001*).
